# Reduced gene dosage of the psychiatric risk gene *Cacna1c* is associated with impairments in hypothalamic-pituitary-adrenal axis activity in rats

**DOI:** 10.1101/2024.08.08.607145

**Authors:** Anna L. Moon, Elle Mawson, Patricia Gasalla Canto, Lawrence Wilkinson, Dominic Dwyer, Kerrie L. Thomas, Jeremy Hall

**Affiliations:** Neuroscience and Mental Health Innovation Institute, School of Medicine, Cardiff University, Cardiff, UK; School of Psychology, Cardiff University, Cardiff, UK; Centre for Neuropsychiatric Genetics and Genomics, School of Medicine, Cardiff University, Cardiff, UK

## Abstract

Common and rare variation in *CACNA1C* gene expression has been consistently associated with neuropsychiatric disorders such as schizophrenia, bipolar disorder, and major depression, however the underlying biological pathways that cause this association have yet to be fully determined. In this study, we present evidence that rats with a reduced gene dosage of *Cacna1c* have increased basal corticosterone levels in the periphery and reduced *Nr3c1* gene expression in the hippocampus and hypothalamus. These results are consistent with an effect of *Cacna1c* dosage on hypothalamus-pituitary-adrenal (HPA) axis function. We also show that the reduction of *Nr3c1* in the hippocampus may be caused by epigenetic modification of exon 1_7_ of *Nr3c1*, including the reduced interaction with the histone modifying markers H3K4me3 and H3K27ac. Heterozygous *Cacna1c* rats additionally show increased anxiety behaviours. These results support an association of *Cacna1c* heterozygosity with the altered activity of the HPA axis and function in the resting state and this may be a predisposing mechanism that contributes to the increased risk of psychiatric disorders with stress.

## Introduction

Genetic variation in *CACNA1C*, the gene that encodes the pore forming subunit of Ca_v_1.2, an L-type voltage-gated calcium channel (L-VGCCs), has been consistently associated with an increased risk for neuropsychiatric disorders, including schizophrenia and bipolar disorder [1–4]. Single nucleotide polymorphisms in intron 3 of this gene have been particularly implicated in heightening risk and have been demonstrated to regulate *CACNA1C* gene expression including in the brain [5–7]. Although the direction of effect has not been consistent across studies [5,6,8,9], there is evidence that *CACNA1C* risk alleles can result in a reduction in Ca_v_1.2 expression especially in the hippocampus [10]. L-VGCCs play a crucial role in cell excitability [11] and are known to mediate neuronal excitation-transcription coupling [12]. Hence, alterations in Ca_v_1.2 expression may have significant implications for neuronal function. Indeed fine-mapping combined with eQTL analysis in the largest Schizophrenia GWAS to date identified *CACNA1C* as a key candidate contributing to the enrichment of gene sets relevant to synaptic function associated with disease [13].

Prolonged, excessive stress and dysregulation of hypothalamic-pituitary-adrenal (HPA) axis has been consistently linked to the development of psychiatric disorders [14–18]. Recent studies have shown that SNPs in *CACNA1C* significantly interact with stress exposure to alter risk for depressive symptoms [19,20] and bipolar disorder [21]. Further, Klaus and colleagues demonstrated that the main risk SNP rs1006737 in *CACNA1C* interacted with early-life stress to determine cortisol-awakening response – which is often used clinically as a read-out of HPA axis activity [22]. These studies indicate a potential moderating role of *CACNA1C* variants on the HPA axis and the association between stressors and risk for psychiatric disorders. Preclinical studies have also revealed a link between *Cacna1c* and the stress system; mice with altered *Cacna1c* expression in the forebrain and nucleus accumbens show increased susceptibility to chronic stress [19,23] and a 5-HT neuron specific knock-out of *Cacna1c* disrupted active stress-coping behaviours [24]. In addition, animal studies show Ca_v_1.2 expression and function to be highly responsive to corticosteroid hormones and stress [25–31]. Thus, the interplay of Ca_v_1.2 and the HPA axis may represent a mechanism for risk of developing neuropsychiatric disorders.

When the body experiences an external or internal stressor the HPA axis is stimulated; the hypothalamus releases corticotropin-releasing hormone (CRH) and arginine vasopressin (AVP) together which instigate the release of adrenocorticotrophic hormone (ACTH) from the pituitary gland. In turn, ACTH stimulates the release of glucocorticoids from the adrenal cortex, namely cortisol in humans or corticosterone in rodents [32]. The production of cortisol/corticosterone forms part of an adaptive stress response regulating a whole range of diverse functions on multiple systems throughout the body including metabolism, cognition and inflammation via the binding of cortisol/corticosterone to low affinity glucocorticoid receptors (GRs) or high affinity mineralocorticoid receptors (MR) in the peripheral tissue and in the brain including the hippocampus, hypothalamus and pituitary through both membrane-bound nongenomic mechanisms to control neuronal activity and nuclear transcriptional activity to regulate gene expression [33]. The binding of cortisol/corticosterone to central GRs also promotes a negative feedback mechanism which inhibits further cortisol production, terminating the acute stress response [34–37]. MRs, which are largely ligand-bound and activated under basal conditions, exert a tonic inhibition of the HPA axis thereby controlling the threshold of reactivity of the system [38–40].

This mechanism confers adaptive advantages when stress is acute and relatively brief via homeostatic and allostatic regulation of metabolic, cognitive, behavioural, neuroimmune and bodily functions appropriate to promoting survival. However, when stress is chronic or unmanageable, the system may become maladaptive and even pathogenic [41]. This leads to a net change in basal cortisol levels. Hypercortisolemia and reduced negative feedback of GCs is associated with depression, general anxiety and panic disorder as well as a number of other psychopathologies [42,43]. Hypocortisolemia and high stress sensitivity has been associated with both traumatic and chronic stress exposure, particularly early-life or childhood trauma experiences, and conditions such as post-traumatic stress disorder, atypical depression, and chronic fatigue syndrome [44–46].

This study sought to directly investigate the hypothesis that variation in *Cacna1c* dosage impacts the HPA-axis using a *Cacna1c* heterozygous rat model). Given the evidence of a reciprocal relationship between activation of the HPA axis and Ca_v_1.2 expression and function after stress, we examined the system under basal, non-stress-associated conditions with the explicit aim of better understanding the causal mechanisms through which genetic variation in *CACNA1C* might contribute to risk for psychiatric disorders.

## Methods

### Cacna1c^+/-^ rats

*Cacna1c* heterozygous rats (*Cacna1c^+/-^*) on a Sprague Dawley background (TGR16930, Horizon, Sage Research Labs, USA) and wild-type littermates were bred in-house and housed in mixed-genotype groups of 2-4. This model has been characterised previously in our lab and others to have 40-50% decrease in *Cacna1c* mRNA and protein levels throughout the brain [10,47]. All animals were housed in standard cages (38cm x 56cm x 22cm) on a 12:12hr light-dark cycle with ad libitum access to food and water unless otherwise stated. Experiments were conducted in accordance with local ethics guidelines, the UK Animals (Scientific Procedures) Act 1986 and the European Communities Council Directive (1986/609/EEC). Gene and hormone expression, and epigenetic modification measures were measures in mPFC, whole hippocampus (dorsal plus ventral) and hypothalamus as indicated from a cohort of all male rats. Further hormone expression assessments and behavioural experiments were conducted on a separate mixed sex cohorts.

### RT-qPCR

RNA was extracted from tissue using the Qiagen RNeasy Kit (Qiagen, UK), following the supplied protocol, using 20-30mg tissue per sample. RNA content was measured on a NanoDrop spectrophotometer to determine concentrations and purity. RNA was DNAse treated (Ambion TURBO DNA-free Kit) before being converted to cDNA. 1.5 µg RNA was added to cDNA conversion (Clontech RNA to cDNA Random Hexamers) tubes and placed in a thermal cycler and heated at 42⁰C for 75 minutes and at 80⁰C for 15 minutes. cDNA was diluted 1:15 in nuclease-free water. Primers for qPCR were designed using the FASTA gene sequences from NCBI (http://ncbi.nlm.nih.gov) and tested for homology using nBLAST before being synthesised by Sigma-Aldrich. Primers (10μM) were validated for single-amplicon specificity using a standard curve analysis. 96-well plates were loaded with 15μl reaction mixture (1:25 primers, 1:3 cDNA and 1:2 SYBR-Green SensiMix (Bioline)). *Gadph* and *Hprt* primers were used as housekeeping controls. Primer sequences were: *Nr3c1 F:* 5’- AGCACAATTACCTTTGTGCTGGA, R: 5’-TTCGATAGCGGCATGCTGGA; *Nr3c2* F: 5’- AATAACGTCCCTCTGCGCTC, R: 5’-GCCTGAAGTGGCATAGCTGA; *Gapdh* F: 5’-TCTCTGCTCCTCCCTGTTCT, R: 5’-TACGGCCAAATCCGTTCACA. *Hprt*: F: 5’-TCCTCCTCAGACCGCTTTTC, R: 5’- ATCACTAATCACGACGCTGGG. qPCR was run as follows: 95⁰C for 10 minutes, 45 cycles of 95⁰C (15s) and 60⁰C (60s), 55⁰C (60s) and 95⁰C (15s). Quantification was done using the comparative Ct method (2^-ΔΔCT^) method to produce fold changes between groups tested.

### ELISA

Samples were collected between 10.00AM and 14.00PM during the rat’s subjective day with collection times randomised across groups. Trunk blood was extracted from rats following sacrifice and transformed to a sterile 2ml lithium heparin Vacuette tube (Grenier Bio-One). Tubes were placed in a centrifuge and span at 4000rpm for 10 minutes at 4⁰C and the separated plasma extracted. Plasma was analysed for corticosterone concentration using the Abcam CORT ELISA Kit (ab108821) according to the manufacturer’s instructions, with a 1:100 plasma dilution. CRH was analysed using CRH ELISA Kit (ABIN2871385, Antibodies Online), according to the supplied protocol.

### ChIP-qPCR

Tissue was dissected into 1mm pieces and treated using the EpiQuik Tissue ChIP Kit (P-2003-2, Epigentek, USA). Tissue was fixed using 1% formaldehyde at room temperature for 5 minutes to cross-link intracellular DNA and protein, washed in ice-cold PBS before the tissue was disrupted by homogenisation. The resulting suspension was spun at 5000rpm for 5 minutes to form a pellet. This pellet was re-suspended in lysis buffer with protease inhibitors and subject to sonication (20 cycles of 30s on, 30s off) using a Diagenode Bioruptor Plus. Cell fragments were centrifuged at 14,000 rpm for 10 minutes at 4⁰C to pellet out cell debris. Supernatant was then added to a 96-well plate with various antibodies: RNA polymerase II (positive control), mouse antibody immunoglobulin G (IgG) (1 mg/ml) (negative control) or antibodies against H3K27ac (ab4729, Abcam) or H3K4me3 (ab8580, Abcam). A 5% input sample, not subject to any antibody treatment, was also taken from the remaining supernatant. Decrosslinking was performed following immunoprecipitation and proteins treated by proteinase K. DNA was eluted and purified using Qiagen QIAquick PCR Purification Kit. Primers for ChIP-qPCR were designed to target the promoter region of target genes using the method detailed above. The promoter region of *Gapdh* was used as a positive control, whilst the promoter region of *Sat2* served as a negative control. Primer sequences were: - *Nr3c1 Exon 1_7_* F: 5’- CTGTAGCCCCTCTGCTAGT, R: 5’- TAGTTTCTCTTCTCCCAGGC; *Nr3c1 (−1kb TSS)* F: 5’- AAGGGTTAGAAGGAATTTGGGGA, R: 5’- TGACGTGCCAGAGCCAATTA: *Gapdh* F: 5’- TCTCTGCTCCTCCCTGTTCT, R: 5’- TACGGCCAAATCCGTTCACA; *Sat2* F: 5’- ACAGCTACTGGAAACGGCTGA, R: 5’- CTCAGGGCTTCTTCACTGATCT. qPCR was performed as described above. ChIP-qPCR data was analysed using the percentage of input method which represents the amount of DNA pulled down by using the antibody of interest relative to the amount of total starting chromatin.

### Pyrosequencing

Genomic DNA was extracted from hippocampal tissue using the DNeasy Blood and Tissue Kit (Qiagen). Unmethylated cytosines were sodium bisulphite converted to uracil using the EpiTect Sulphite Kit (Qiagen), while methylated cytosines remained unchanged. Primers were designed to target *Nr3c1* exon 1_7_ using PyroMark Assay Design software and synthesised by Sigma Aldrich (HPLC purified). The reverse primer was 5’ end biotin labelled, to allow for binding to streptavidin beads. Primer sequences for *Nr3c1 Exon 1_7_* were thus F: 5’- TTGTTATTTTAGGGGGTTTTGGTT, R: 5’-[Biotin] AAAAAAACCCAATTTCTTTAATTTCTCTTC. Bisulphite converted DNA was subject to two rounds of PCR (PyroMark PCR Kit, Qiagen) to obtain the amplicon of interest. This biotinylated PCR product was bound to Streptavidin Sepharose HRP beads (Cytiva) and agitated on a shaker for 10 minutes at 1500rpm. A handheld vacuum plate (Qiagen) was used to isolate the bead-amplicon by denaturing the dsDNA strand to obtain a single biotinylated strand. This strand was then annealed to the sequencing primer (5’- GGGGGTTTTGGTTGT) by heating to 80⁰C for two minutes. A Q96 cartridge (Qiagen) was prepared for pyrosequencing. Both cartridge and plate containing the DNA were run on the PyroMark machine to obtain methylation percentages for the region.

### Behavioural measures of anxiety

In a separate cohort of rats behavioural testing was conducted between the hours of 10:00 and 16:00, with a pseudo random distribution of testing for rats of different genotypes across the test day. Rats were habituated to the test rooms for 30 min prior to testing. All assays involved individual testing of rats and apparatus was cleaned thoroughly with a 70% ethanol solution between subjects. Testing in the elevated plus maze (EPM) occurred 3 days after testing in the open field (OF). Both tests are widely used paradigms for assessing innate anxiety indexed by the competition between the exploration of novel contexts and the aversion to open, brightly lit environments or heights [48–50].

The OF consisted of an opaque white plastic box (100 x 100 x 40 cm high) illuminated at 30-35 lx. During testing, animals were placed into the centre of the box and its location monitored for 10 min.

Elevated plus maze consisted in two open arms and two closed arms made of black plexiglass walls and wood black floor, elevated by 70cm and in low light conditions (30-35 lx). Each open and closed arm measured 50 x 10 cm, with closed arms surrounded by 10 cm high walls. Animals were place in the centre of the EPM facing an open arm and were recorder for 5 min. EPM was surrounded by a dark blue curtain from ceiling to floor in order to avoid any other external context stimulation. Equivalent arms were arranged opposite one another. Rats were placed at the enclosed end of a closed arm and allowed to freely explore for 5 min.

Data for the OF and EPM and were collected using EthoVision XT software (Noldus Information Technology, Netherlands) using a digital camera mounted above the centre of each piece of apparatus. Each rat was tracked (12 frames/s) for location of its greater body-proportion in specific zones within each apparatus. For the OF, two virtual zones of equal area were created corresponding to the 50% inner or central zone and the 50% outer ring zone closest to the arena walls. For the EPM, data from the open pair of arms were combined to generate a single duration in open arm value.

### Statistics

Data was analysed using *t*-tests to assess whether there were differences between *Cacna1c^+/-^* genotype in outcome measures. Linear regression was used to assess the effect of sex and genotype as well as their possible interaction when both sexes were included in analyses. Two-sided tests were used apart from instances in which results were replicated in a separate cohort with an *a priori* directional hypothesis. All models were checked for normality of residuals and homogeneity of variance by visual inspection of QQ plots and histograms of residuals, and dependent variables were transformed as indicated if required, and the transformation used is specified in each case. All data was analysed using RStudio v2024.04.1.

## Results

### Cacna1c hemizygosity is associated with decreased GR expression in the hypothalamus and hippocampus

*Nr3c1* and *Nr3c2* encode the glucocorticoid and mineralocorticoid receptors respectively, which play a key role in the HPA-axis stress response network. To assess the effect of reduced *Cacna1c* gene dosage on HPA functioning, the effect of genotype of the heterozygous *Cacna1c* rat model on *Nr3c1* and *Nr3c2* gene expression within the brain was measured. In the hippocampus, a stress-sensitive region with a high proportion of *Nr3c1* and *Nr3c2,* a reduction in *Nr3c1* was seen in *Cacna1c* HET rats compared to WT (*t*_(24)_ = −2.654, *p* = 0.014; square root transformed) (Figure 1A). Within the hypothalamus, a key part of the HPA axis, a similar reduction was observed (*t* _(13)_ = −2.486, *p* = 0.027; square root transformed) (Figure 1A), whereas in the PFC, no change was found (*t*_(21)_ = −0.504, *p* = 0.621; log transformed) (Figure 1A). No difference was observed in *Nr3c2* in any region (Hippocampus: *t*_(21)_ = −1.430, *p* = 0.168 (log transformed), Hypothalamus: *t*_(11)_ = −0.805, *p* = 0.438 (log transformed), PFC: *t*_(14)_ = 0.005, *p* = 0.944 (inverse square transformed)) (Figure 1B). mRNA expression data were log transformed. Note that the selective decrease in *Nr3c1* expression in the hippocampus and hypothalamus was seen in two separate cohorts of male HETs and WTs (*data not shown*) and therefore data were combined. The reproducibility of this finding highlights the robustness of the effect of *Cacna1c* gene dose on *Nr3c1* expression in these brain regions.

**Figure 1:**
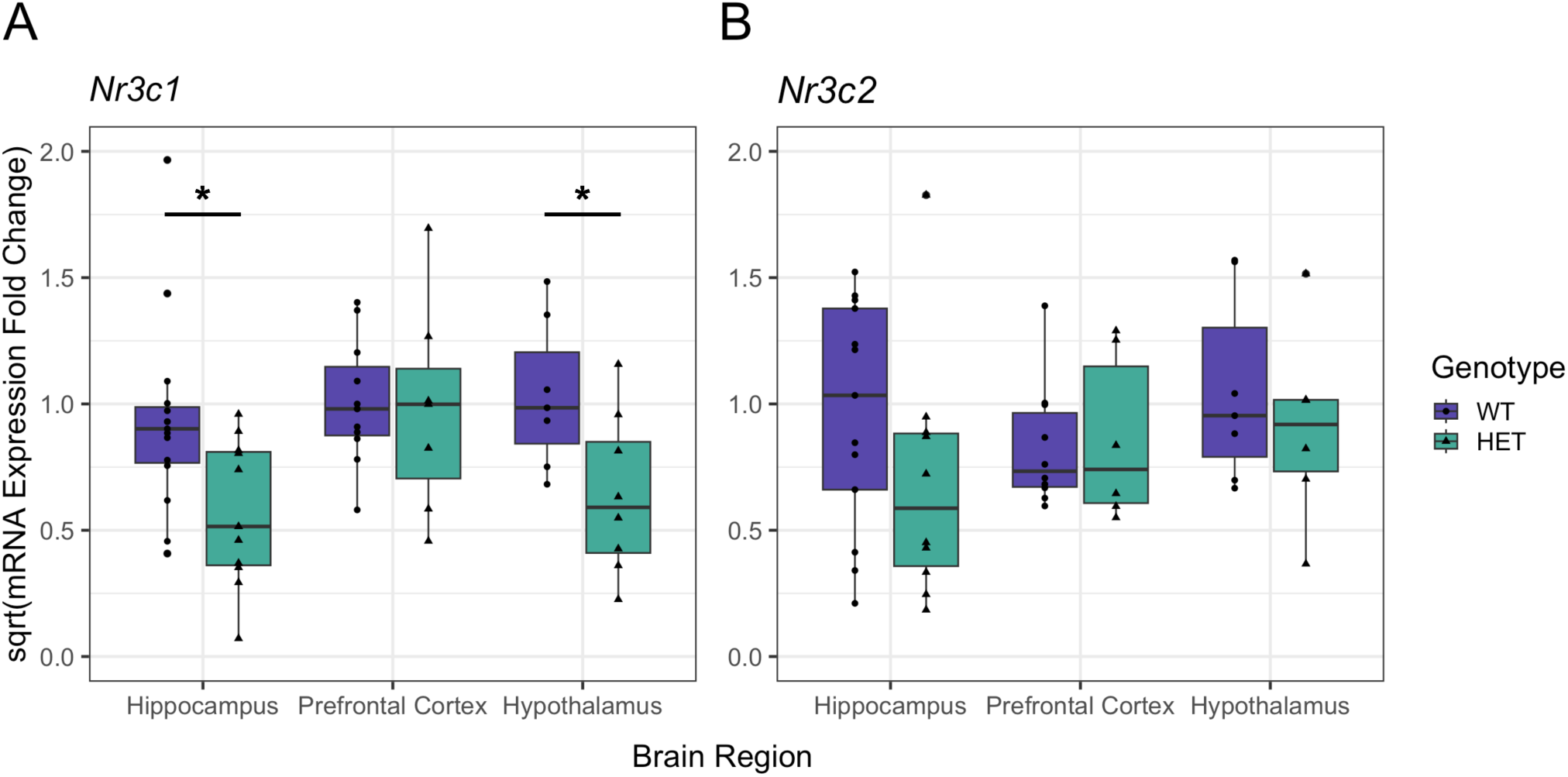
A. *Cacna1c^+/-^* rats (HET) have reduced *Nr3c1* gene expression in the hippocampus and the hypothalamus compared to wild-types (WT) B. There are no differences in *Nr3c2* expression in any brain region tested. Data are shown as mean ± SE. Results were considered significant if *p* < 0.05 (*),. *Nr3c1*: Hippocampus, HET n = 11, WT n = 15; PFC, HET n = 7, WT n = 11; Hypo HET n = 8, WT n = 8. *Nr3c2*: Hippocampus, HET n = 9, WT n = 12; PFC, HET n = 6, WT n = 10; Hypo HET n = 6, WT n = 7.

### Epigenetic changes in exon1_7_ of Nr3c1 in the hippocampus of heterozygous Cacan1c rats

#### DNA methylation

The decrease in *Nr3c1* in *Cacna1c^+/-^* rats may be due to epigenetic effects. The *NR3C1* gene contains several alternative non-coding exon 1 variants that contain sites of epigenetic modification that can regulate *NR3C1* gene transcription. Focus has been on exon 1_7_ in rats, which has been demonstrated to have an epigenome that is susceptible to environmental influences such as early life maternal care, resulting particularly in changes in DNA methylation at CpG site 16 containing the 5’ binding site for NGFIA and altered hippocampal GR levels [51,52]. Furthermore, in humans DNA methylation in the analogous 1F CpG cluster region has been associated with psychopathologyl [53]. We therefore designed primers to explore DNA methylation in this region including the CgG site 16 within exon 1_7_. In the hippocampus we found no significant differences in methylation of any of the investigated CpG sites between the male HETs and WT with the exception of decreased methylation in CpG site 14 observed in *Cacna1c^+/-^* rats compared with WT animals (*t*_(22)_ = −3.006, *p* = 0.007) (Figure 2A, Supplementary Table 1)

**Figure 2:**
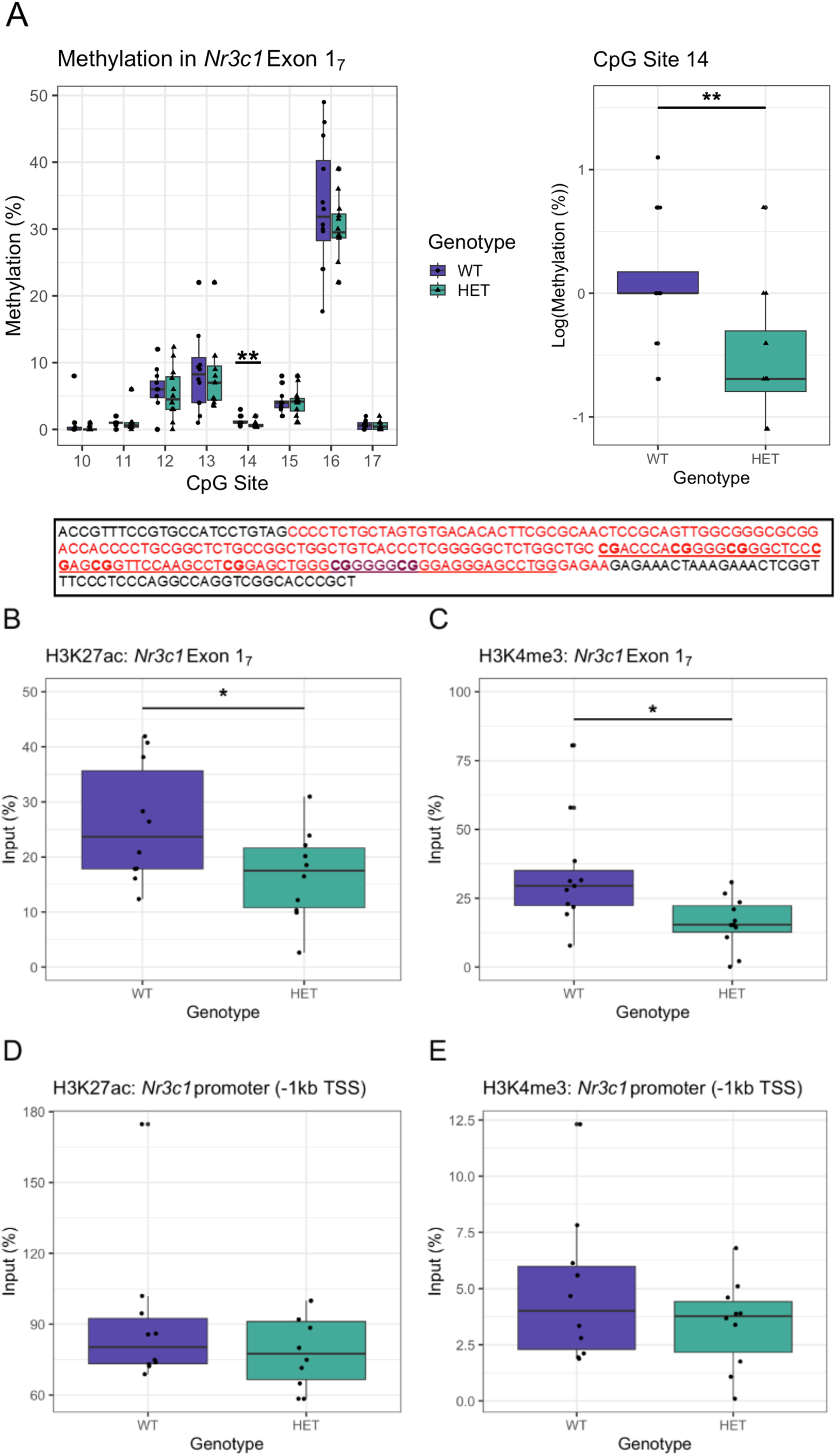
A. *Top*: The % of methylation at each targeted CpG site in the hippocampus of WT and *Cacna1c^+/^-* rats is displayed (*Top Left*) with no significant differences between the genotypes, with the exception of CpG 14 (*Top Right*). The highest % methylation was within at the 5’ CpG site (CpG_16_) within the NGFIA binding site. *Bottom*: Schematic showing the genetic sequence that indicates Exon 1_7_ within the promoter region of *Nr3c1* in the rat [85]. The red region indicates the sequence of Exon 1_7_, with the NGFIA binding site contained within, highlighted in purple. The sequence analysed by this study is underlined and the CpG dinucleotides investigated are in bold (corresponding to CpGs 10*-*17[86]. B-E: *Cacna1c^+/-^* rats show reduced DNA interacting with histone modification markers of active transcription H3K4me3 and H3K27ac within the exon 1_7_ region (B, C). No differences were seen in the region closer to the transcription start site (E, E). n = 10 per group, * = p < 0.05.

#### Histone modifications

As there was no change of direct DNA methylation within seven of the eight targeted CpG sites in exon 1_7_, we utilised ChIP-qPCR to investigate potential histone modifications that correlated with the expression of *Nr3c1* in the hippocampus of hemizygous rats. Histone modifications in the exon 1_7_ GR promotor region associated with altered GR expression in the hippocampus have been reported [51,52,54]. Tri-methylation of H3K4 and acetylation of H3K27 has been shown to be a markers of active gene transcription [55,56]. Using primers to target *Nr3c1* exon 1_7_ (Supplementary Table 5). We observed that in *Cacna1c^+/-^* compared to WT rats there was a reduction in H3K27ac (*t*_(18)_ = −2.169, *p* = 0.044) (Figure 2B) and H3K4me3 (*t*_(20)_ = −2.722, *p* = 0.013; square root transformed) (Figure 2C) interactions within the *Nr3c1* exon 1_7_ region. Using primers designed to target a region −1kb upstream of the TSS within the *Nr3c1* promoter region we showed that there was no significant genotype difference in DNA interacting with H3K27ac (*t*_(18)_ = 1.111, *p* = 0.281; inverse transformed) (Figure 2D) or H3K4me3 (*t*_(19)_ = −1.190, *p* = 0.249; square root transformed) (Figure 2E). These results suggest that reduced *Nr3c1* expression in the hippocampus of *Cacna1c^+^*^/-^ rats could be at least partly driven by altered histone modification within the promoter of *Nr3c1* within exon 1_7._ Despite a degree of spatial selectivity in altered H3K27ac or H3K4me3 levels in the GR promotor, there was no difference in mRNA expression of exon 1_7_ itself (Supplementary Figure 1) in heterozygote rats.

### Cacna1c heterozygosity increases peripheral corticosterone levels

Corticosterone and corticotrophin releasing hormone (CRH) concentrations were measured in blood plasma in an initial all-male cohort of *Cacna1c^+/-^* rats. There was no difference in peripheral CRH (*t*_(22)_ = 0.011, *p* = 0.991; log transformed) (Figure 3A), but there was a significant increase in peripheral corticosterone in *Cacna1c* HET animals compared to WT (*t*_(36)_ = 2.078, *p* = 0.045; log transformed) (Figure 3B).

**Figure 3:**
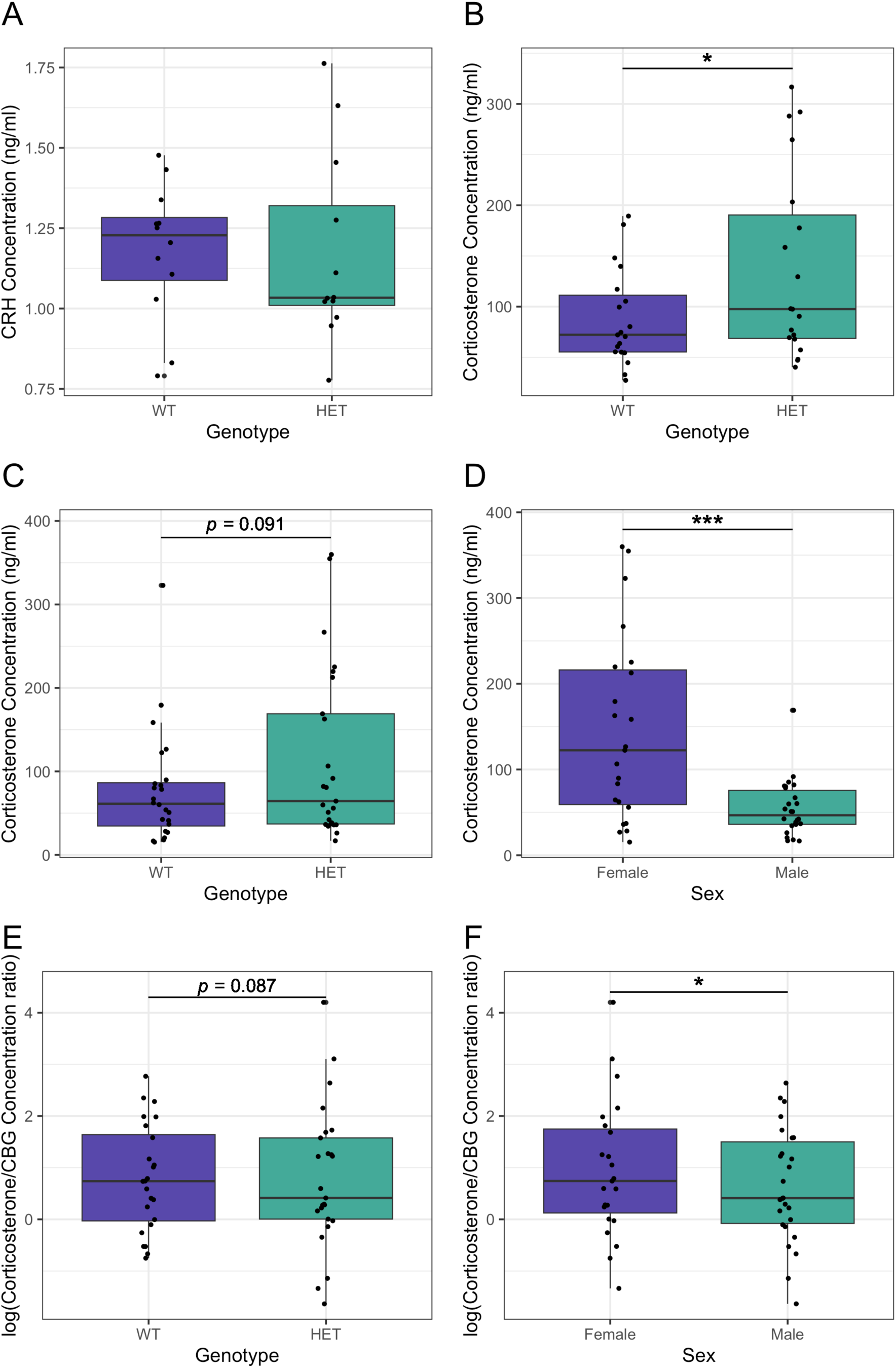
A. Male *Cacna1c^+/-^* rats (HET) have higher circulating corticosterone levels than WT males (n = 19 per group). B. Male *Cacna1c^+/-^* rats have similar CRH hormone levels to male WT (n = 12 per group). C. In a separate mixed sex cohort (n: Male WT = 13 Male HET = 13 Female WT = 11 and Female HET = 12), *Cacna1c^+/-^* rats showed an increased peripheral corticosterone concentration compared to WT. D. A profound sex difference in corticosterone levels was also observed, with females showing higher levels than males. E. There was a trend towards an increase in the ratio of peripheral corticosterone/CBG of HETs (*p* = 0.0866). F. Corticosterone/CBG was higher in females than males. Measures from individual rats are shown as black dots. Bold and narrow horizontal bars denote mean ± SEM, respectively. * p < 0.05, *** p < 0.001.

To assess whether this genotype difference reflected bioavailable corticosterone levels, and to determine whether the effect was found in both sexes, corticosterone, and corticosteroid-binding globulin (CBG) were subsequently measured in a mixed-sex cohort. A sex by genotype interaction was not significant (*t*_(46)_ = −0.950, *p* = 0.347; log transformed) and moreover the AIC score was improved by removing this term from the regression model. Elevated plasma corticosterone levels in *Cacna1c^+/-^*rats compared to WT was seen in the mixed sex cohort (*t*_(47)_ = 1.427, *p* = 0.080, one-tailed and log transformed, Figure 3C), In addition, females were shown to have higher corticosterone concentration compared with males (*t*_(47)_ = 3.749, *p* < 0.001; log transformed) (Figure 3D).

When corticosterone travels through the blood, it is bound to CBG which provides protection from degradation but also when bound to CBG, corticosterone is unable to act on target tissues [57]. Hence, assessing whether there are differences in the ratio of corticosterone to CBG is important to better understand how changes in corticosterone might impact upon HPA-axis targets. A higher corticosterone to CBG ratio would indicate more bioavailable corticosterone. There was a trend towards a higher corticosterone/CBG ratio in *Cacna1c^+/-^* rats compared to WT (*t*_(46)_ = 1.753, *p* = 0.087; one-tailed) (Figure 3E). An increased corticosterone/CBG ratio was also observed in females compared to males (*t*_(46)_ = 2.151, *p* = 0.0370) (Figure 3F). No interaction was observed between sex and genotype (*t*_(46)_ = −1.527, *p* = 0.134). Together this data suggests that the increase in circulating corticosterone in *Cacna1c* heterozygous rats is reflected in bioavailable levels in both sexes.

### Increased anxiety-associated behavioural responses in Cacna1c heterozygous rats

To determine whether the elevated circulating corticosterone levels we observed in the *Cacna1c^+/-^*rats was correlated with increased anxiety like behaviours [58] we assessed emotional reactivity as the time spent in the central zone of an open field and in the open arms of the EPM.

*Cacna1c^+/-^* rats spent less time in the inner 50% of the OF arena than WT (*t*_(53)_ = −2.108, *p* = 0.040; square root transformed, Figure 4A). There was no difference between the sexes (*t*_(53)_ = −1.335, *p* = 0.188; square root transformed) and no interaction between sex and genotype (*t*_(53)_ = 1.525, *p* = 0.133; square root transformed) on the time spent in the inner 50% zone of the OF. There were no differences in locomotor activity between the genotypes or the sexes analysed as total distance travelled (Supplementary Figure 2A).

**Figure 4.**
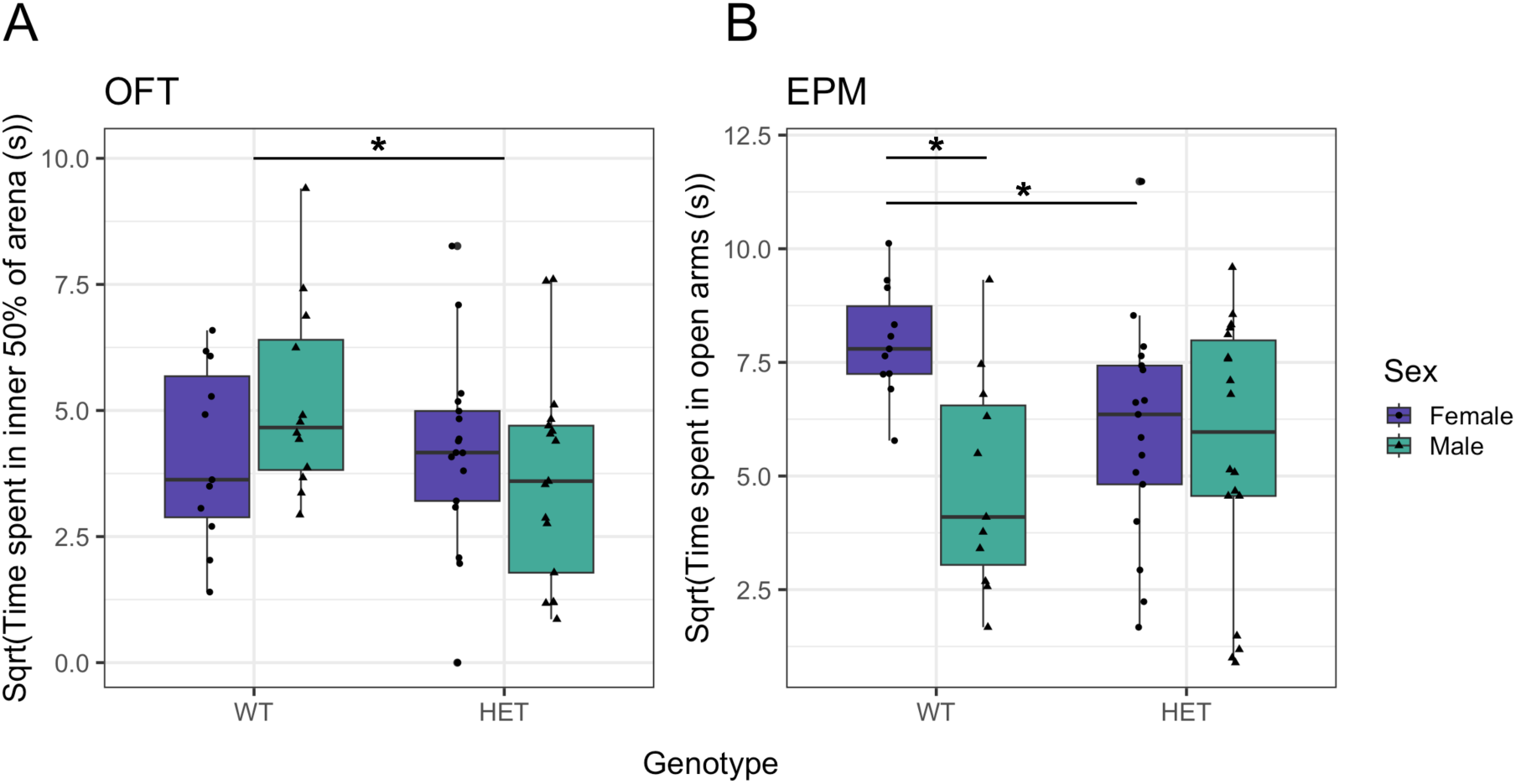
A. There is a reduction in time spent in the inner 50% of an OF arena in *Cacna1c^+/^-* rats (HET) compared to WT animals B. In the same cohort, there were no differences of genotype on the time spent in the open arms of the EPM. However, female rats spent more time in the open arms compared to males and there was a significant interaction between sex and genotype (t(53) = −2.040, p = 0.0463), where female *Cacna1c^+/^-* rats showed a reduced time in open arms compared to female WT (t(26) = - 2.457, p = 0.021). Dependent variables were square root transformed in to adhere to model assumptions. Measures from individual rats are shown as black dots. Horizontal bars denote the median. * p < 0.05. OFT: Male HET n = 17; Male WT n = 12; Female HET n = 17; Female WT n = 11. EPM: Male HET n = 18; Male WT n = 12; Female HET n = 17, Female WT n = 11.

In the EPM there was no differences in the time spent in the open arms (*t*_(53)_ = −0.772, *p* = 0.444; square root transformed, Figure 6A) in the *Cacna1c^+/-^*. Nevertheless, there was an interaction between genotype and sex in time spent in the open arms (*t*_(53)_ = −2.040, *p* = 0.046; square root transformed), with *Cacan1c*^+/-^ females showing decreased time in open arms compared to WT females. This was reflected by decreased time spent in the open arms in HET females compared to WT females (*t*_(26)_ = - 2.457, *p* = 0.021; square root transformed), with no difference between genotypes found in males (*t*_(27)_ = 0.689, *p* = 0.497; square root transformed). Females spent significantly more time in the open arms than males (*t*_(53)_ = 3.007, *p* = 0.004; square root transformed), concomitant with females showing increased (*t*_(53)_ = −2.002, *p* = 0.050) locomotor activity compared to males (Supplementary Figure 2B).

## Discussion

This study aimed to assess the effects of reduced *Cacna1c* gene dosage on HPA-axis functioning using a heterozygous *Cacna1c* rat (HET) model. We observed a decreased level of *Nr3c1* mRNA encoding the GR in the hippocampus and the hypothalamus of *Cacna1c^+/-^* rats. The reduction of *Nr3c1* expression in the hippocampus correlated with reduced H3K4me3 and H3K27ac, histone modifications associated with activating gene expression, in exon 1_7_ of *Nr3c1*. The selective decrease in GR and not MR expression in hippocampus and hypothalamus coupled with the increased plasma CORT levels in the absence of changes in CRH levels measured under baseline conditions are suggestive of GR resistance and altered HPA feedback in the HETs [33]. Thus, constitutive knockdown of *Cacna1c* expression results in the development of an HPA axis that may function differently to that normally. Increased innate anxiety responses were also observed in this *Cacna1c^+/-^* rat model.

Alongside the effects on basal circulating corticosterone levels, we also show a reduction of *Nr3c1* gene expression in *Cacna1c^+/-^* rats in the hippocampus and hypothalamic regions but not the PFC. This indicates a degree of regional selectivity of *Cacna1c* hemizygosity on HPA axis function. As such it is interesting to note that GR activity in the hippocampus and PVN is central to the negative feedback required for allostatic adaptive regulation of the HPA axis after stress [33]. DNA methylation, particularly at the CpG site 16 of *Nr3c1*, is inversely associated with NGFIA transactivation of GR expression and expression of exon 1_7_-containing GR transcripts in the hippocampus [52,59]. We saw no evidence of altered DNA methylation at CpG_16_ of the exon 1_7_ in the *Cacna1c^+/^* rats and it is then perhaps not surprising that we saw unaltered expression of exon 1_7_-containing GR transcripts. Nevertheless, we measured epigenetic changes in exon 1_7_ *Nr3c1* in the *Cacna1c^+/^* rats, including decreased CpG_14_ methylation and decreased H3K4me3 and H3K27ac, modifications that are correlated with transcriptional activation and repression respectively. Whilst there is evidence of epigenetic changes within *Nr3c1* associated with *Cacna1c* hemizygosity, the full epigenetic signature that results in reduced GR expression in the hypothalamus and hippocampus associated with *Cacna1c* haploinsufficiency remains to be determined.

DNA demethylation in at *Nr3c1* CpG_16_ and active histone marks including H3K27ac at the exon 1_7_ site have been positively correlated with GR expression and inversely correlated with depressive-like behaviour [51,52,54]. Notably, the dynamic relationship between these functional epigenetic signatures and behaviour have been observed in studies following a period of early life stress. In this regard our results showing no changes in CpG_16_ exon 1_7_ methylation status and exon 1_7_ transcript expression under basal, non-stressed conditions in the *Cacna1c^+/-^* rats are concordant with the CpG_16_ methylation at exon 1_7_ of *Nr3c1* being mechanistically linked to stress-associated functional adaptations and psychopathy’s [53]. Nevertheless, histone modifications within exon 1_7_ in *Cacna1c^+/-^*rats provides evidence for the susceptibility of this region to epigenetic alteration following either environmental stress or genetic influences.

The allosteric load model conceptualises that acute stress may be adaptively advantageous in instigating biological coping mechanisms, but that chronic, excessive, or uncontrollable stress leads to a transition to a defective state that is maladaptive [41]. Glucocorticoids and their receptors play a central role in this model, typically with a U-shaped function such that high and low levels are associated with suppressed or non-engaged synaptic and neuroplasticity mechanisms required for adaptation, respectively [60]. Building on this model, Chrousos and colleagues [61] suggest that this latter vulnerable state associated with chronic hyperactivation of the HPA, elevated circulating CORT and decreased negative feedback by GR can transition over time to a self-preserving downregulation of the HPA axis characterised by reduced CORT production. Key variables that affect the transition along the adaptive-to-maladaptive continuum include the chronicity and intensity of the HPA activation by stressors, and also the developmental timing of the stress exposure [61]. In this light, the hypercortisolism and GR resistance we observe in the *Cacna1c^+/-^* rats at may represent a model of heightened vulnerability to stress and consequential altered cognitive, behavioural, metabolic and neuroimmune functions associated with a range of psychiatric disorders.

The *Cacna1c^+/-^* rats show increased anxiety-like behaviours compared to WT. Our observations in the rat model are consistent with those seen in the haploinsuffiency mouse model [62,63]. The change in anxiety-like behaviour was not confounded by genotype effects on motoric behaviour, similar to that reported for haploinsufficient mice at least in males [62,64]. Increased anxiety-like behaviour measured in the *Cacna1c^+/-^* rats was most markedly shown in the OF test where decreased dwell time in the more innately aversive, open central regions of the walled arena is seen irrespective of sex. When using the EPM, increased anxiety as measured at time in the open arms was seen in female *Cacna1c^+/-^* rats only. These observations may reflect that the OF and EPM tasks differentially engage motivational components of exploration (fear vs. curiosity that together mediate risk taking in open spaces) in so much as the OF is forced exploration of an open environment [65]. Our data in rats together with previous studies in mice [63] suggest that females appear to be more sensitive to the effects of Ca_v_1.2 reduction on behavioural measures of anxiety than males indicating sex-specific effects on underlying motivational processes. The sex-dependent sensitivity on anxiety-like behaviour in *Cacna1c^+/-^* rodents concurs with the observation that women with *CACNA1C* risk variants are more prone to mood disorders and anxiety than men [63,66].

It is important to note that reports of altered anxiety-like behaviour in rodent models of *Cacna1c* deletion may also depend on the cell type or brain regions targeted. Like global *Cacna1c* haploinsufficiency models as discussed above, in models where there is a conditional KO in principle excitatory neurones, particularly in PFC [67], or D1R expressing excitatory neurones [64], increased anxiety-associated behaviours are seen. However, models with excitatory and inhibitory neuronal Ca_v_1.2cKO express normal anxiety-like behaviour [68,69]. Studies in cell-restrictive models of *Cacna1c* deletion are informative for determining the locus of anxiety dysregulation, or indeed other functional impacts of altered Ca_v_1.2 levels, and the underpinning neural mechanisms (e.g. in considering E/I balance). This knowledge will be required for the development of targeted therapies informed by arguably the more disease-relevant models of *Cacna1c* haploinsufficiency that likely impact Ca_v_1.2 levels in glial as well as neuronal populations [70] in a regionally selective manner [10].

The increased plasma corticosterone levels in the *Cacna1c^+/-^*rats may at least in part underlie the increased anxiety response given that corticosterone treatment and stress activation of the HPA is associated with the development of anxiety-like behaviours in female and male rodents [58,71–74]. The decrease in GR expression we measured in the hippocampus and hypothalamus may also contribute to the mechanism. As discussed above, reduced GR deletion and subsequent reduced negative feedback to the HPA is associated with increased circulating CORT. In addition, there is evidence that deletion of GR forebrain excitatory cells including the hippocampus, but excluding the PVN, is associated with increased anxiety-like behaviour and hypercortisolism [75]. Of note we also measured no changes in the expression of the MR in hippocampus, which have been associated with a permissive role in CORT regulation of anxiety-like behaviour [76]. Together the data suggest that the elevated MR/GR ratio in hippocampus and perhaps other regions of the neural circuits may underly elevated innate anxiety we measured in the *Cacna1c^+/-^*rats.

We observed higher CORT levels in females compared to which is consistent with previous observations [77,78] and higher bioavailable CORT (higher CORT/CBG). Therefore, contrary to what we observed, a sex difference in anxiety-like behaviour may have been predicted because of the tight association between circulating CORT levels and anxiety. In addition to limiting CORT bioavailability, CBG plays an essential role in regulating CORT release and emotional reactivity and is under the regulatory control of gonadal estrogen [79]. It is possible that differences in the regulation of CBG-associated cellular functions between males and females may account for the similar expression of anxiety-like behaviour in both sexes in the face of differing circulating CORT levels. Accounting for estrous stage in future studies will provide a more complete understanding of the relationship of sex to the behavioural manifestation of anxiety in the *Cacna1c^+/-^*rats.

Our data suggest that the psychiatric risk rat model of *Cacna1c* haploinsuffiency presents with an altered HPA axis in adulthood as evidenced by increased baseline plasma CORT levels, lowered GR expression (and thus evidence of an elevated MR/GR ratio) in the hypothalamus and hippocampus, and associated increases in innate anxiety-like behaviour. This phenotype suggests a rat model displaying an altered vulnerability to stress [80]. It will be interesting in future studies explore the behaviour and physiological consequences of stress in this model, in particular after early-life stress which has been tightly associated with altered HPA axis function and psychiatric illness [46,81,82]. It is important that these studies are conducted in females and males not least because of the sex differences in the development and regulation of the HPA axis [83] and the importance such studies for the development of treatments for psychiatric disease in men and women [84].

## Supporting information

Moon Supplemental table and figures

## Acknowledgements

We thank J. Carter for technical support. This work was funded by MRC grant no. MR/R011397/1 and legacy funds, with further support from the Hodge Centre for Translational Neurosciences

## Notes

### Competing Interest Statement

The authors have declared no competing interest.

