## Supplementary material for "Reduced gene dosage of the psychiatric risk gene *Cacna1c* is associated with impairments in hypothalamic-pituitary-adrenal axis activity in rats": Moon Supplemental table and figures

**Supplementary Table: Statistical reports for pyrosequencing**

| **CpG** | $t_{(22)}$ **value** | ***p* value** |
|---|---|---|
| 10 | -1.022 | 0.318 |
| 11 | -0.029 | 0.978 |
| 12 | -0.199 | 0.844 |
| 13 | 0.024 | 0.981 |
| 14 | -3.006 | 0.007 |
| 15 | -0.137 | 0.892 |
| 16 | -1.015 | 0.321 |
| 17 | <0.001 | >0.999 |

**Supplemental Figures**

**Supplemental Figure 1**

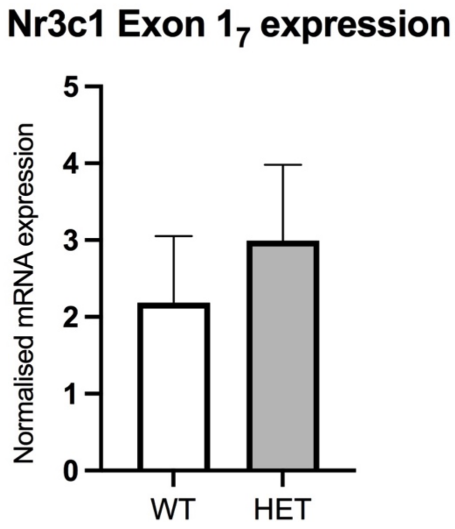


Supplementary Figure 1: No differences were seen in the expression *of Nr3c1 exon* $1_7$ in the hippocampus between wild-type and *Cacna1c* hemizygous rats ($p$ = 0.555, n = 8 per group) as measured by qPCR. Expression levels were normalised *Gapdh* and *Hprt* expression. Data are shown as mean ± SE.

**Supplemental Figure 2**

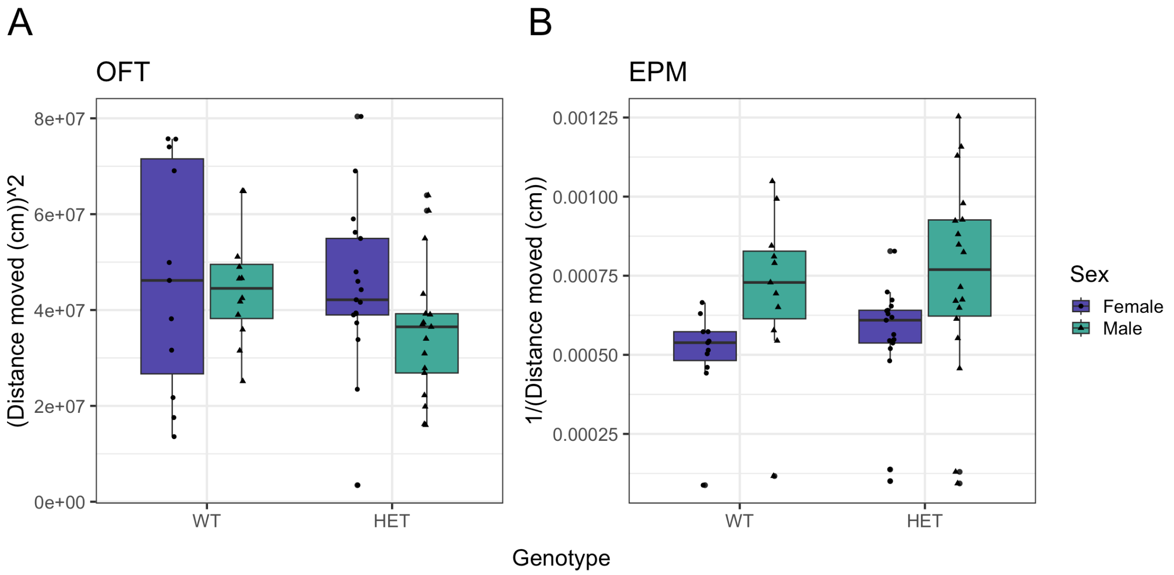


Supplementary Figure 2: Figure 2: A. In the OF, there we no effect of *Cacna1c* haploinsufficiency or sex on the locomotor activity as measured by distance travelled in cm (genotype: $t_{(53)}$ = -1.440, $p$ = 0.156), sex: ($t_{(53)}$ = 0.248, $p$ = 0.805, genotype x sex: $t_{(53)}$ = 0.775, $p$ = 0.442). B. In the EPM, while there was no effect of *Cacan1c* hemizygosity on activity as measured by total distance travelled ($t_{(53)}$ = 0.431, $p$ = 0.668), females travelled more than males ($t_{(53)}$ = -2.002, $p$ = 0.050). There were no genotype x sex interactions ($t_{(53)}$ = 0.089, $p$ = 0.930). Dependent variables were square transformed in A and inverse transformed in B to adhere to model assumptions. Measures from individual rats are shown as black dots. Horizontal bars denote the median. OFT: Male HET n = 17; Male WT n = 12; Female HET n = 17; Female WT n = 11. EPM: Male HET n = 18; Male WT n = 12; Female HET n = 17, Female WT n = 11.
